# Measuring stimulus-related redundant and synergistic functional connectivity with single cell resolution in auditory cortex

**DOI:** 10.1101/2023.06.19.545531

**Authors:** Loren Koçillari, Marco Celotto, Nikolas A. Francis, Shoutik Mukherjee, Behtash Babadi, Patrick O. Kanold, Stefano Panzeri

## Abstract

Measures of functional connectivity have played a central role in advancing our understanding of how information is communicated within the brain. Traditionally, these studies have focused on identifying redundant functional connectivity, which involves determining when activity is similar across different sites. However, recent research has highlighted the potential importance of also identifying synergistic connectivity—that is, connectivity that gives rise to information not contained in either site alone. Here, we measured redundant and synergistic functional connectivity with individual-neuron resolution in the primary auditory cortex of the mouse during a perceptual task. Specifically, we identified pairs of neurons that exhibited directed functional connectivity between them, as measured using Granger Causality. We then used Partial Information Decomposition to quantify the amount of redundant and synergystic information carried by these neurons about auditory stimuli. Our findings revealed that functionally connected pairs carry proportionally more redundancy and less synergy than unconnected pairs, suggesting that their functional connectivity is primarily redundant in nature. Furthermore, we observe that the proportion of redundancy is higher for correct than for incorrect behavioral choices, supporting the notion that redundant connectivity is beneficial for behavior.

## 1 Introduction

Functional connectivity (FC) has emerged as a mainstream concept and a fundamental tool to understand how networks in the brain communicate, and how functional interactions between networks or between neurons shape the dynamics and functions of the brain [1, 2, 3, 4, 5]. Most of these traditional measures of FC focused on redundant connectivity, by measuring (for example, through linear (cross-)correlations) the similarity of activity at different sites. However, recent studies have begun to highlight the importance of another type of FC — synergistic connectivity. This type of connectivity focuses on how variations of the interaction between activity at different sites create information that is not present at each site alone [6, 7, 4, 8]. While the presence and merits of redundant connectivity have been extensively documented [1, 3, 9], it remains unclear whether synergistic interactions are prominent and how they contribute to functions.

An additional question regards the spatial scale at which both redundant and synergistic interactions are expressed. Most studies of FC investigated it at a coarse scale, such as that obtained with fMRI or EEG [10, 11, 12, 13, 14]. The organization of FC at the smaller spatial scale of population recordings with single-cell resolution is less understood.

In this study, we address some of these open questions regarding synergistic and redundant FC. First, we address their relationship with respect to a widely used FC measure, Granger Causality (GC) [15, 16], between the activities of different neurons. This measure of FC is interesting because, unlike simple measures of FC based on cross-correlation, it considers not only the similarity of activity but also the strength and directionality of information transmission. GC can, in principle, capture redundant FC because the process of transmission causes sharing of information between the sending and the receiving site. However, it can also correspond to synergistic FC. For example, if transmission varies across sensory stimuli, FC can create sensory information not available in each node individually. Second, we use precise information theoretic measures to quantify redundancy and synergy related to the encoding of important sensory variables (in this case, features of auditory stimuli). These measures, based on the theory of Partial Information Decomposition (PID) [17], have the advantage of separating redundancy from synergy, something that simpler measures [18] used in recent studies [19, 20] cannot do. Third, we study synergy and redundancy with neuron resolution, using the primary auditory cortex (A1) of the mouse brain as an experimental model. Fourth, we explore the potential impact of synergy and redundancy on sensory processing by studying how they vary between cases of correct and incorrect perceptual discrimination.

## 2 Experimental task and single neuron stimulus information

To investigate the relationship between FC and the presence of synergistic and redundant information with single-neuron resolution, we focused on the activity of the mouse primary auditory cortex during a sound discrimination task. We reanalyzed a previously published dataset [20] in which the activity of several tens to a few hundreds of neurons was recorded simultaneously using *in vivo* 2P calcium imaging (Ca^2+^) from A1 L2/3 neurons in transgenic CBA*×*Thy1-GCaMP6s F1 mice during a pure-tone discrimination task (Fig. 1A).

**Figure 1:**
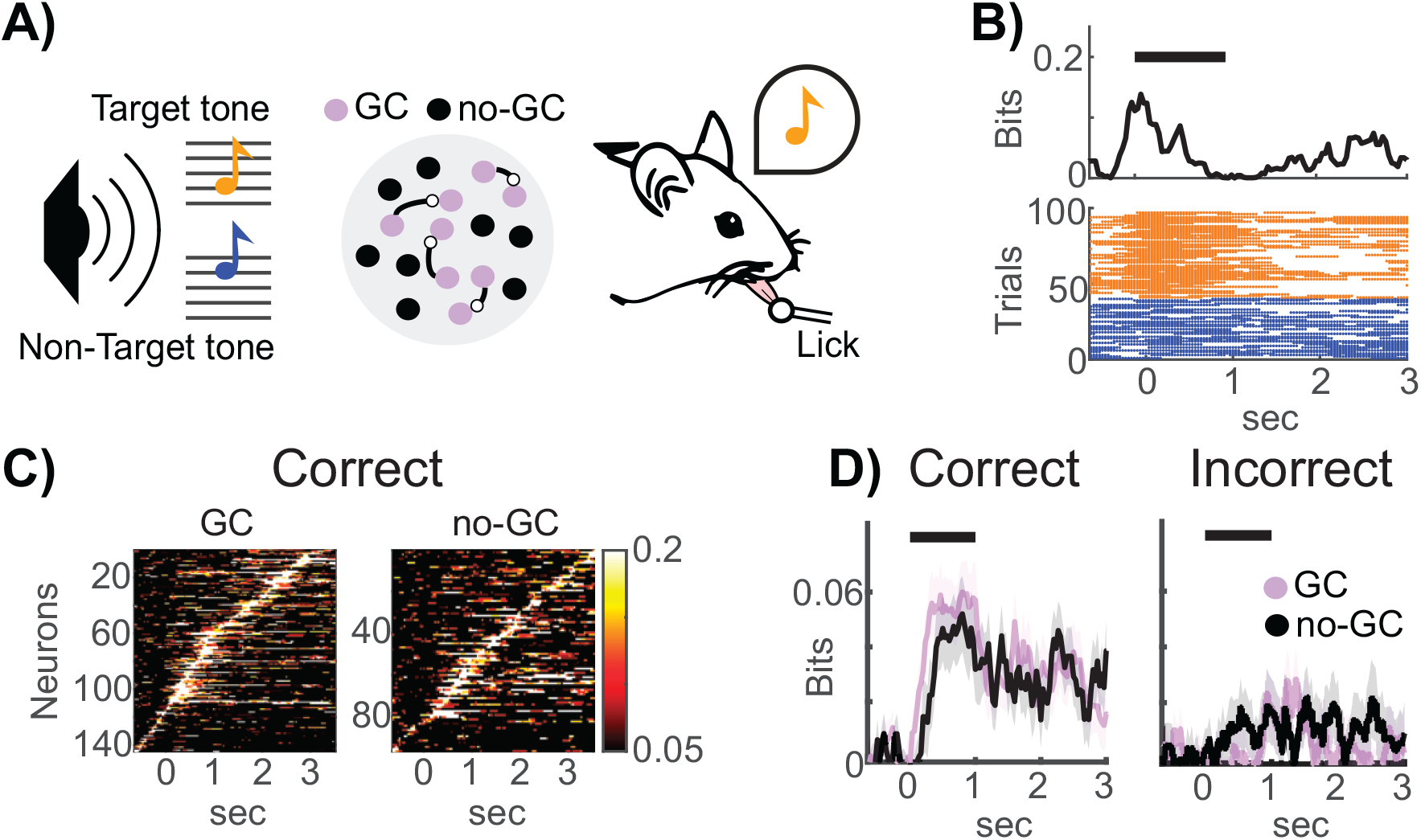
Stimulus information in mouse auditory cortex during a tone discrimination task. A) Mice performed a go/no-go tone discrimination task while the activity of A1 L2/3 neurons were monitored with two-photon calcium imaging. In response to a target tone (low-frequency, in orange) mice had to lick a waterspout and not to lick for non-target tones (high-frequency, in blue). Granger Causality analysis revealed sparsely connected networks of cells in A1 L2/3 [20]. We classified neurons as GC (purple) and no-GC (black) depending on whether or not they formed a GC link. B) Example of the stimulus information time-course for a single neuron. We computed the time-resolved stimulus information as the mutual information (top plot) between the auditory stimuli (low-/high-frequency tones) and the spike activity across trials. In the raster plot (bottom plot) we color-coded and sorted the spiking activity based on the tone that occurred across trials. C) Stimulus information time-course for GC neurons (left map) and no-GC neurons (right map) in correct trials only. We then sorted the peaks of stimulus information for each neuron to tile the trial time. D) Population-averaged stimulus information for GC (in purple) and no-GC (black) neurons in case of correct and incorrect trials separately.

The experimental task was structured as follows. After a pre-stimulus interval of 1 second, head-fixed mice were exposed to either a low frequency (7 or 9.9kHz) or a high frequency (14 or 19.8kHz) tone for a period of 1 second. Mice were trained to report their perception of the sound stimulus by their behavioral choice, which consisted of licking a waterspout in the post-stimulus interval (0.5 to 3 seconds from stimulus onset) after hearing a low-frequency tone (target tones) and holding still after hearing high-frequency tones (non-target tones). Calcium imaging was used to continuously acquire the fluorescence signals from individual A1 L2/3 neurons during the task with a sampling frequency of 30Hz.

We used Shannon mutual information [21, 22] to compute the stimulus information carried by each neuron about the stimulus category (low vs high-frequency tones) in each imaging time frame (Fig. 1B, top plot). Stimulus information is defined as follows:

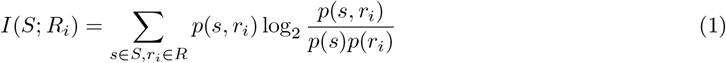

where *i* indexes the neurons and *p*(*s, r*_*i*_) denotes the joint probability of observing in a given trial the activity *r*_*i*_ of neuron *i* and the value *s* of the stimulus variable *S. p*(*r*_*i*_) = Σ_*s*_ *p*(*s, r*_*i*_) and *p*(*s*) = Σ_*r*_ *p*(*s, r*_*i*_) are the marginal probabilities. We estimated *p*(*s, r*_*i*_) using the direct method [23].

The activity *r*_*i*_ of neuron *i* was inferred following the same approach described in [20]. Briefly, we first deconvolved the single-trial calcium fluorescence traces of each neuron to infer the spiking activity (Fig. 1B, bottom plot). We then used a sliding window approach (a sliding window of 10 time frames with time-steps of 1 time frame) to binarize the spiking activity into 0 and 1, where 1 denotes spiking activity higher than zero. We then computed the time-resolved stimulus information on these neural responses. Finally, we subtracted the average stimulus information computed in the pre-stimulus interval from the stimulus information time-courses, which enabled us to remove the bias in the information that occurred for the limited number of trials [24].

Following our previous study [20], we first analyzed the entire dataset (2792 neurons recorded from 34 sessions) to identify those neurons that carried significant task-related information. Neurons were defined as carrying task-related information if they carried statistically significant stimulus information (defined as in Eq 1 above), significant choice information (defined as in Eq 1 above but replacing the stimulus presented in the given trial with the behavioral choice of the animal in the trial), and intersection information, defined as the amount of sensory information encoded in neural activity that is used to inform the behavioral choice [25] and computed using PID [26]. The statistical significance of each information measure was computed using a non-parametric permutation test at *p <* 0.1. The requirement of all three non-independent tests being satisfied simultaneously was empirically estimated, resulting in a false discovery rate of 1% [20]. We found a subset of 475/2790 neurons that transiently and sequentially carried significant task-relevant information[20]. Using methods described in [16, 20] we next performed a Granger causality (GC) analysis on the selected subset of neurons. We picked the 20 neurons with the peak intersection information with the shortest latencies in 12 of 34 sessions and found that they formed sparse functional networks that transmitted redundant task-relevant information across the trial time (Fig. 1A)[20]. Of these 240 neurons, 144 formed GC connections with other neurons in the network and were termed GC neurons hereafter. The remaining 96 neurons, which did not form GC connections with other neurons, were termed no-GC neurons hereafter.

We used information-theoretic measures to map the stimulus information dynamics of individual neurons. We first considered information in trials in which the mouse made correct perceptual discriminations to understand whether GC neurons and no-GC neurons encoded information differently during correct behavior. The stimulus information time-courses, plotted in (Fig. 1C) after sorting neurons by their peak information timing, showed sequential information coding across the population in both GC and no-GC neurons. Neurons in both classes had similar amounts of information at peak, the main difference being that GC-connected neurons had peak information earlier in the trial (during the stimulus presentation), while no-GC neurons carried information later in the trial (after stimulus presentation) (Fig. 1C). The sequential nature of their activation suggests that information is represented throughout the trial only at the population level, motivating our later information analyses at the neural population level.

To investigate what aspects of neural activity may be key for performing correct perceptual judgements, we assessed how information about the auditory stimulus category was encoded in trials in which the animal judged the stimulus either correctly or incorrectly. For simplicity, we averaged the stimulus information across neurons (Fig. 1D). Importantly, we found that in incorrect trials, the stimulus information dropped for both populations across the entire trial time, suggesting that the stimulus information that both GC and no-GC neurons encode is used for the behavioral choice. Importantly, all information quantities computed during correct discriminations were calculated on random subsets of correct trials with the same size as the number of incorrect trials in the same session. With this equalized sub-sampling, we could fairly compare the amount of information encoded in correct and the incorrect trials, thereby controlling for potential artefacts introduced by different amounts of limited-sampling bias [24].

## 3 PID for measuring synergy and redundancy

The above analysis considered the single-neuron correlates of correct and incorrect perceptual discriminations with the directed FC. We next asked how these properties relate to the emergent properties of population coding. This requires computing stimulus information from more than one neuron.

As in our previous study [20], we estimated the total stimulus information that was jointly carried by pairs of neurons, following a time-lagged approach (Fig. 2A). We first identified for each neuron the peak times of task-related information, i.e., the time frames when intersection information time-courses peaked. We then computed the time-lagged stimulus information carried jointly by the activity of each pair of neurons as follows:

**Figure 2:**
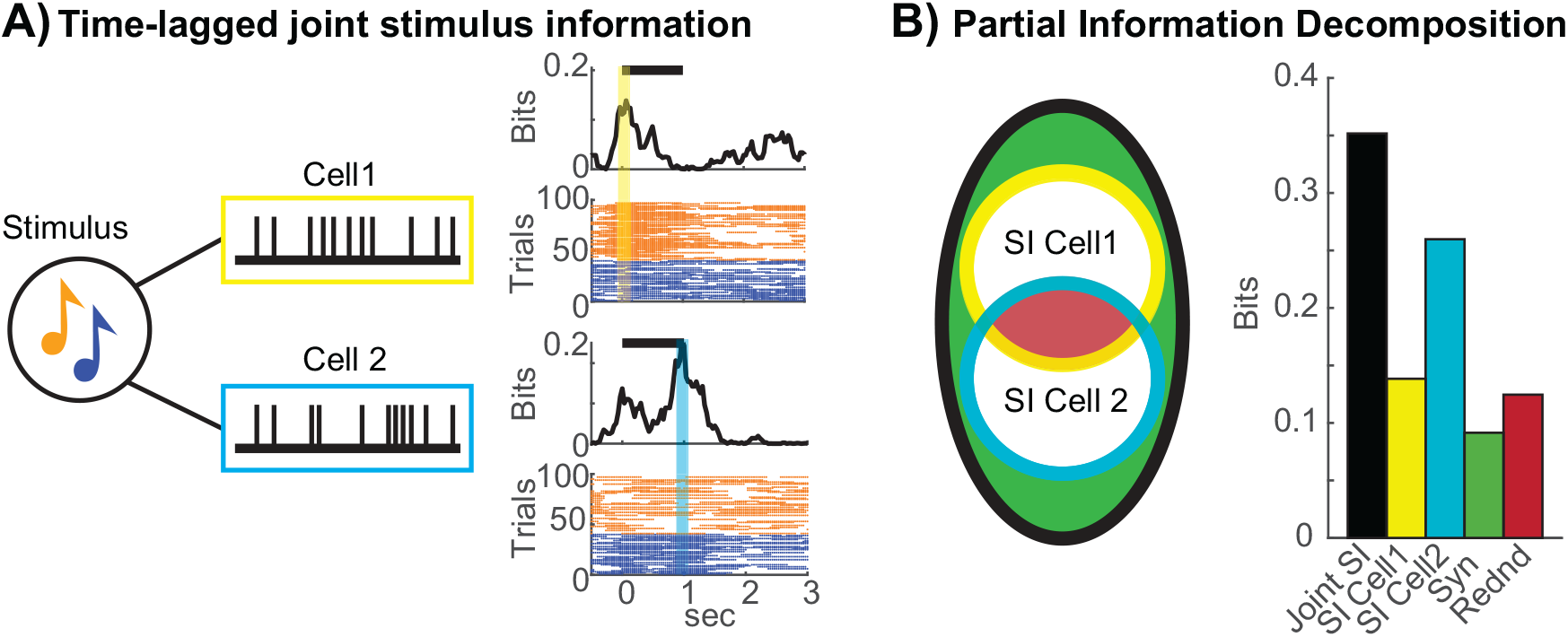
Partial information decomposition of the time-lagged joint stimulus information. A) Time-lagged joint stimulus information is defined as the mutual information that the neural responses of two cells at their information time peaks (yellow and cyan vertical bars in the raster plots) jointly carry about the stimulus. We colour-coded and sorted trials in the raster plots based on the stimulus category (low vs high-frequency tones). B) The joint stimulus information can be decomposed into the non-negative components of synergy, redundancy, and the stimulus information of each cell (Venn diagram on the left). The bar plot on the right shows the amount of the joint stimulus information for the pair of cells in panel A) (black bar), stimulus information of both cells (yellow and cyan colors), the synergistic (green bar), and redundant (red bar) components.

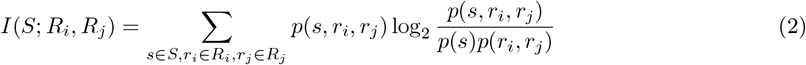

where *p*(*s, r*_*i*_, *r*_*j*_) denotes the probability of simultaneously observing in the same trial the value *s* of the stimulus category and the joint responses *r*_*i*_ and *r*_*j*_ of neurons *i* and *j* at the time frames of their peak of task-related information. In our previous work [20], we investigated the nature of redundant and synergistic interactions in pairs of neurons by computing the so-called co-information [27, 28], defined as the difference between the total stimulus information that was jointly carried by both neurons (Eq 2 above) and the sum of the information carried by the single neurons individually (Eq 1 above):

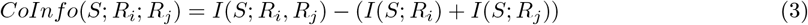

If *CoInfo*(*S*; *R*_*i*_; *R*_*j*_) is positive, then the pair of neurons carries overall more information than expected by summing the information carried by the neurons individually. Thus, a positive sign of this quantity is interpreted as predominant synergy, and a negative sign as predominant redundancy. However, it has been shown that co-information conflates two non-negative pieces of information which properly and separately quantify synergy and redundancy [17]. Indeed, there could be cases when co-information is low, but synergy and redundancy are both high [29]. In simple terms, redundancy (the area in red in the Venn diagram in Fig. 2B) quantifies the amount of information that both neurons carry independently about the stimulus, while synergy (the area in green in the Venn diagram in Fig. 2B) is the amount of information that can be accessed when observing both neuronal responses simultaneously but is not carried individually by any neuron.

To determine the specific contributions of synergy and redundancy to the total joint information, we used the formalism of PID [17]. PID allows breaking down the joint mutual information that two or more source variables carry about a target variable into non-negative and well-interpretable pieces of information. An important insight coming from this decomposition is that *CoInfo* is the difference between two distinct pieces of information that can be quantified separately:

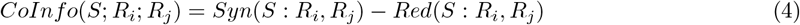

where *Red*(*S* : *R*_*i*_, *R*_*j*_) is the redundant information which is present in both neuron *R*_*i*_ and neuron *R*_*j*_, and *Syn*(*S* : *R*_*i*_, *R*_*j*_) is the synergistic information carried only by the joint response of the two neurons. To compute *Red*(*S* : *R*_*i*_, *R*_*j*_) and *Syn*(*S* : *R*_*i*_, *R*_*j*_) we used the definition provided by [30], where the two terms are computed by solving a constrained convex optimization problem in the space of the trivariate probability distributions *Q*(*S, R*_*i*_, *R*_*j*_). To numerically solve this optimization problem, we used the *BROJA_2PID* python package [31]. In this way, we decomposed the joint stimulus information into the synergy, redundancy, and individual neuron stimulus information.

## 4 Measuring stimulus-related synergy and redundancy in auditory cortex

In our previous study [20] we found an overall redundancy of task-related information in correct trials and an overall synergy in incorrect trials for both pairs of neurons having a GC link and pairs of neurons not having a GC link, suggesting that redundancy is relevant for the behavioural choice. Following [20], we labelled neuronal pairs as GC-connected if they shared at least one GC link and as GC-unconnected otherwise. Since we employed the conflated measure of co-information in [20], the above levels of redundancy in correct trials could arise in distinct scenarios: redundancy is higher in correct rather than in incorrect trials, synergy is lower in correct than in incorrect trials, or a combination of the two. To separate the amounts of stimulus-related synergistic and redundant interactions, we thus employed the formalism of PID. We next performed the stimulus-related PID analysis in the two separate groups of GC-connected and GC-unconnected pairs of neurons in correct and incorrect trials.

We first performed the PID analysis in correct trials for both the GC-connected and GC-unconnected pairs of neurons (Fig. 3A). Again, we randomly sub-sampled the correct trials to equalize the number of incorrect trials (average over 100 repetitions). The joint stimulus information had comparable values (0.386*±*0.002bits, 0.402*±*0.012bits for no-GC and GC pairs respectively) in both populations. However, GC-connected pairs had higher levels of redundancy (0.121*±*0.006bits) compared to the GC-unconnected ones (0.105 *±*0.001bits), while they had similar amounts of synergy (0.097*±*0.001bits, 0.093*±*0.003bits for GC-unconnected and GC-connected pairs respectively) (Fig. 3A). The difference between synergy and redundancy, i.e., the co-information (Eq 4), showed a prevalence of redundant information in both populations, but the GC-connected pairs were more redundant (*−*0.027*±*0.006bits) than GC-unconnected pairs (*−*0.007*±*0.001bits).

**Figure 3:**
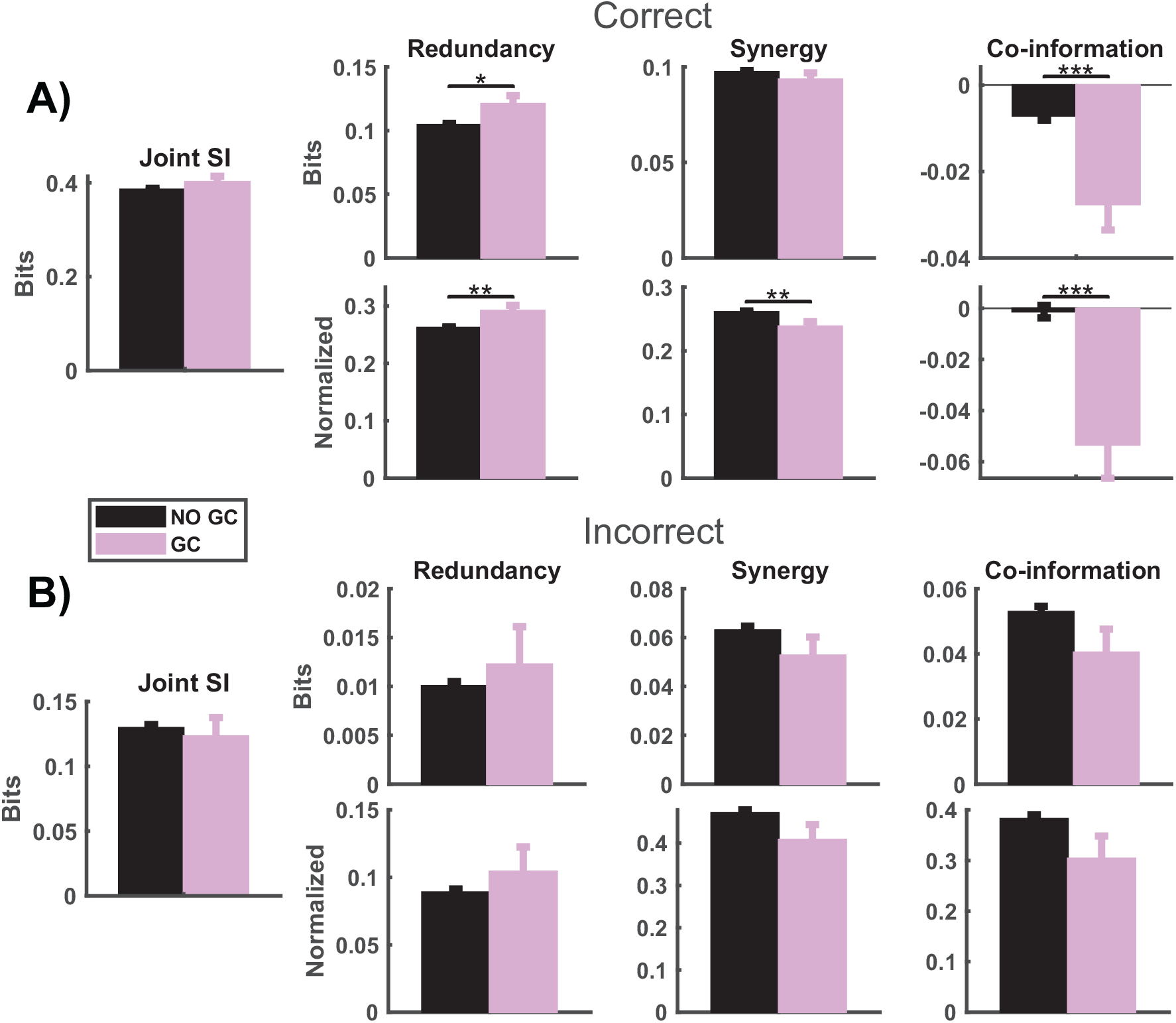
Redundancy and synergy in auditory cortex. A) Redundancy and synergy PID computed in correct trials. From left to right: time-lagged joint stimulus information (SI), redundancy, synergy, and co-information for GC-connected (purple) and GC-unconnected pairs of neurons (black). For synergy, redundancy and co-information, the top plots show values in bits and the bottom plots show values normalised by the joint stimulus information. B) As in panel A), but in the case of incorrect trials. Bar plots show mean ± SEM across pairs. Statistics were made with a Wilcoxon rank-sum test (*p < 0.05, **p < 0.01, ***p < 0.001).

We next quantified the fraction of redundancy and synergy by normalizing each term with respect to the total joint mutual information. We found that GC-connected pairs had proportionally more redundancy and less synergy (red= 0.292*±*0.01, syn= 0.239*±*0.007), compared to GC-unconnected ones (red= 0.262*±*0.001,syn= 0.261*±*0.001) (Fig. 3A). Finally, the normalized co-information showed that GC-connected pairs neurons had much higher redundancy (0.053 0.013) than GC-unconnected pairs (0.001 0.002). Our results suggest that GC-connected pairs of neurons have more redundant than synergistic functional connections.

Next, we investigated whether larger amounts of redundancy and lower amounts of synergy could be beneficial for task performance and behavioral accuracy. We computed the PID in incorrect trials (Fig. 3B). The joint stimulus information in incorrect trials was only about ∼30% of what it was in correct trials (0.130*±*0.002bits, 0.123*±*0.014bits for no-GC and GC pairs respectively). Redundancy in incorrect trials had a value of (0.01*±*0.001bits, 0.012*±*0.004bits for no-GC and GC pairs respectively), which is proportionally ten times smaller than that of correct trials, while synergy dropped to (0.063 *±*0.002bits, 0.053*±*0.007bits for no-GC and GC pairs respectively), proportionally only half of that in correct trials. Co-information showed positive values, i.e. more synergy than redundancy, in both GC-unconnected and GC-connected pairs (0.053*±*0.002bits, 0.04*±*0.007bits). Fractional redundancy was just ∼10% of the total information, whereas fractional synergy was ∼45% of it (Fig. 3B). Moreover, we did not find significant differences in the normalized co-information between GC-unconnected and GC-connected pairs on incorrect trials (0.382*±*0.008, 0.304*±*0.044). Our results suggest that only the redundant FC associated with GC links is useful for correct sensory discriminations.

## 5 Discussion

In this study, we teased apart the relationship between FC and stimulus-related synergy and redundancy with single-neuron resolution in the auditory mouse cortex during a perceptual discrimination task. We deliberately considered one specific, widely-used type of directed FC measure, Granger Causality. Unlike other measures such as the Pearson correlation between the activity of two neurons, Granger Causality can in principle be related to redundancy and synergy. Our findings revealed that Granger FC between A1 L2/3 neurons was accompanied by proportionally higher levels of redundancy and lower levels of synergy compared to pairs of neurons that were not linked with a Granger FC. Moreover, we found that the levels of redundancy were higher in both populations in correct behavioral choices compared to incorrect ones. Our results suggest that both synergy and redundancy coexist across the population, regardless of whether or not they are Granger connected and of whether or not the mouse makes correct or incorrect perceptual discriminations. However, redundancy becomes proportionally more prominent, and synergy less prominent, when Granger FC is present and during correct behavior. Overall, these results suggest that FC creates prevalent redundancy of sensory information across neurons, and that this redundancy is beneficial for correct sensory judgements. The advantages of redundancy for perceptual discrimination found here could arise from multiple contributions. One well-documented advantage regards the integration of information across sites [32]. Another one could result in advantages in terms of information transmission and readout. Indeed, while redundancy limits the amount of encoded information [33], it has benefits in terms of improving the propagation of information between pre- and post-synaptic neurons [9, 4]. Together with those reported in previous studies [9, 20, 4], our results suggest that the optimal trade-off between the advantages and disadvantages of redundancy results in an overall advantage of having some degree of redundancy to secure safe downstream information transmission.

Our finding confirms previous reports of the presence of significant synergy between the activity of neurons or networks [19, 8]. However, our finding of decreased synergy during correct perceptual discrimination suggests that the potential advantages of synergy in terms of higher levels of encoding of sensory information do not necessarily or directly translate into advantages for sensory discrimination. One possibility is that these types of synergistic information may be more difficult to read out, as it would require more complicated decoders that may be beyond the capabilities of some downstream neural circuits. However, given that presence of synergy has been well documented, another possibility, to be explored in future studies, is that synergy may not be needed for the simple perceptual tasks we consider but that it could become more important for more complex behaviors.

From the theoretical perspective, previous studies that investigated synergy and redundancy between neurons or networks employed a measure of co-information which conflates synergy with redundancy, measuring only their net effect [18, 20]. With respect to these studies, we made the advance of using a more refined measure that teased apart redundancy from synergy, which allowed us the important step forward of being able to measure separately their relationship with both FC and the accuracy of behavior. With respect to other studies considering redundancy and synergy, but not relating it to information content about variables of cognitive interest [8], we made progress by measuring redundancy and synergy of information about variables, such as sensory stimuli, which have a well-defined meaning and role in terms of perceptual functions. We hope that our work contributes to creating a neuroinformatics framework that can help researchers to study the patterns of synergy and redundancy about external stimuli and pinpoint their contribution to behavior and functions.

In conclusion, our findings suggest that correct behavior is associated with a pervasive presence of redundant information in functionally connected neural networks. Further research is needed to better understand the contributions of synergy and redundancy in different contexts.

